# Roles of *PKCdelta* in Photoperiod and Circadian Regulations in *Drosophila melanogaster*

**DOI:** 10.1101/2021.09.29.462474

**Authors:** Harjit Khaira, Somya Patro, Kwangwon Lee

## Abstract

Many organisms are known to regulate seasonal behaviors and physiological processes in response to day length changes through photoperiodism. Extreme changes in photoperiods have detrimental effects on human health, which can impair development and serve as the origin of adult diseases. Since the seminal work by Bünning in 1936, there are studies supporting the view that organisms can measure the day length through an endogenous 24-hour cellular circadian clock. However, the mechanisms involved in measuring seasonal or day-length changes are not understood. In the current study, we performed a genome-wide association study (GWAS) on photoperiodism using the Drosophila Genetic Reference Panel. The GWAS identified 4 top candidate genes responsible for photoperiodic regulations. The knockout mutants of the four candidate genes (*Protein Kinase C delta* (*Pkcdelta*), *Glucuronyltransferase-P* (*GlcAT-P*), *Brain-specific homeobox* (*Bsh*), and *Diuretic hormone 31 Receptor* (*Dh31-R1*)) were analyzed for their photoperiod and circadian period phenotypes. *PKCdelta* and *GlcAT-P* mutants show a significantly different photoperiod response compared to that of the wild type strain, and also had an altered circadian period phenotype. Further molecular characterization revealed two independent mutant alleles of *PKCdelta* with a defective catalytic domain had distinct photoperiod responses. Taken these data together, we concluded that there is overlap between the circadian clock and photoperiodic regulations in *Drosophila*, and *PKCdelta* is a component that is involved in both circadian and photoperiodic regulations. By identifying novel molecular components of photoperiod, the current study provides new insights into the genetic mechanisms of determining the seasonal changes.

**Author Summary:** Extreme changes in photoperiods have detrimental effects on human health, which can impair development and serve as the origin of adult diseases. The molecular and genetic mechanisms of how an organism interprets and adapts to seasonal environmental changes are not well understood. Drosophila Genetic Reference Panel (DGRP) is a community resource of 205 inbred lines created for studying population genomics and quantitative traits. Using DGRP we performed a genome-wide association study (GWAS) to find potential photoperiod candidate genes. *PKCdelta*, a candidate gene from the GWAS study was identified to have both photoperiod and circadian effects. This data supports the view that circadian clock and photoperiodism have a shared regulatory circuit. Our study sheds light onto potential genes that could be further studied to characterize the mechanisms of photoperiodism, and its relationship to the circadian clock.

## Introduction

Most organisms experience predictable environmental changes; daily, tidal, lunar, and seasonal. These changes mean different availability of resources and stressors at predictable time intervals. It is crucial for an organism to prepare for the incoming season for its survival in nature. Thus, it is not surprising that organisms have evolved time-measuring cellular devices. Among the known biological rhythms, seasonal rhythm plays a major role in many biological processes such as growth, development, reproduction, migration, and dormancy. A robust response to these rhythms can increase the survival success of an organism (1). However, little is known about the molecular and genetic mechanism of how an organism interprets and adapts to seasonal environmental changes.

Photoperiodism is the physiological response of an organism to changes in day length. Photoperiod is one of the most important factors that control seasonal breeding. Few species show extreme behaviors like hibernation while many species synchronize breeding and birth of their offspring to the most advantageous time of the year for better survival. Based on the gestation period, photoperiodic species can be categorized as long day (LD) or short day (SD) breeders. Animals like sheep and goats with long gestation periods begin to breed as day length decreases, while animals like mice and hamsters breed as day length increases (2). In humans there are some psychological and psychiatric consequences in response to the photoperiodic changes. There also is seasonality in birth rates among humans and onset of certain diseases like measles in the population (3). Patients with bipolar disorders have more depressive episodes during the late autumn and early winter, while there is an increase of manic episodes during spring and summer (4). In flowering plants, some are flowering at specific season of the year, and thus grouped as SD or LD flowering plants. The molecular and genetic pathways of photoperiodism have been studied in plants (5). It is also known that the circadian clock controls the activity of CONSTANS (CO), which is a key component of the photoperiod pathway. CO activates the transcription of *FLOWERING LOCUS T* (*FT*) which the ultimately leads to flowering in plants (6).

Photoperiodic responses are well documented in insects (1). Although *Drosophila melanogaster* is successful model species for many biological processes, the species was regarded as less optimal for studying photoperiodism due do a less pronounced photoperiod response (7). Most of the studies of photoperiodism in insects utilized diapause as a phenotype. Diapause occurs when female insect ovaries shrink due to photoperiod changes as an indicator of photoperiodic phenotype. On the contrary to the concern in using *D. meanogaster* for studying photoperiodism, it also has been shown that *D. melanogaster* has a robust photoperiod response under semi-natural or SD conditions (8,9).

One compelling hypothesis regarding the seasonal rhythm is that an organism can ‘interpret’ the season by measuring the day length. A crucial assumption is that an organism knows the season by measuring photoperiods is due to the presence of an endogenous cellular daily clock, called the circadian clock. Circadian clocks are endogenous timekeeping networks that drive 24hr circadian rhythms. Bünning reported that the rate of leaf movement speed changes during different seasons and proposed that the circadian clock controlled this via photoperiodism (10). Since then, a number of studies supported Bunning’s hypothesis in certain organisms while rejecting his hypothesis in others. Studies on aphids and spider mites conclude that their photoperiod mechanisms do not seem to depend on endogenous 24-h rhythms (11). Mutation of the classical clock gene *period* (*per*) still allowed *D. melanogaster* to distinguish between SD and LD. It was concluded that the clock gene *per* cannot be involved in night-length measurement mechanism (12). However, another study in *Drosophila* suggests that the circadian clock is necessary for a photoperiodic response, as arrhythmic clock mutants *per^01^*, *tim^01^*, and *Clk^JRK^* have altered photoperiodic responses from wild-type *Drosophila* (13). A recent study established that known clock protein TIMELESS (TIM) modulates the levels of EYES ABSENT (EYA) protein. EYA is involved in measuring photoperiodic and temperature changes to trigger physiological response such as diapause (14). For the last few decades, the genetic and molecular mechanisms of circadian clocks have been characterized in many organisms (15,16). However, it is still not well understood how the circadian clock is involved in measuring photoperiodism or day length changes.

There are at least two different hypotheses on how circadian clocks are involved in measuring seasonal changes: the external coincidence model and the internal coincidence model. In the external coincidence model, the photoperiodic response can be generated by the circadian clock that is sensitive to light at certain phases (17). If the light signal is detected at this sensitive phase, then the photoperiodic response will be displayed such as flowering in plants. Light can entrain the clock and set the phase of photoperiodic response like flowering, but light can also directly induce flowering. In the internal coincidence model, two (or more) rhythms are produced by the clock. Under the internal coincidence model, responses, such as flowering, are only generated in conditions that bring both rhythms in phase with each other (17). Various studies show one or the other hypotheses at work in a range of organisms. In *Arabidopsis*, external coincidence models can explain long day-specific dusk *FT* (a florigen gene) expression (18). Studies using the parasitic wasp *Nasonia vitripennis* support the internal coincidence model. Using the Nanda-Hamner protocol it was found that *N.vitripennis* exhibit two peaks in diapause, a dawn and dusk peak, which is suggestive of two circadian oscillators (19). In some bacterial species the hourglass model is suggested to be responsible for timekeeping (20). When bacteria exposed to more regular environmental conditions such as consistent light-dark cycles or constant photoperiod, hourglass model is at play. However, when the environmental conditions vary such as inconsistent light-dark cycles can lead to a sustained oscillator (20).

In our current study, we aim to better understand the genetic mechanisms of photoperiodism and the roles of circadian clocks in photoperiodism using the eukaryotic model organism *D. melanogaster*. *Drosophila* Genetic Reference Panel (DGRP) is a community resource of 205 inbred lines created from 40 wild type parents for studying population genomics and quantitative traits (21). Because the DGRP is inbred to homozygosity, it is possible to identify SNP variants responsible for the trait of interest. DGRP is a fully sequenced population with annotated high-quality SNPs, indels, and chromosomal inversions (21,22). DGRP has been used to study various complex traits such as starvation resistance, startle response, chill coma recovery (CCR) time, and circadian behavior (21,23).

Although diapause has been the phenotype of choice in characterizing the photoperiodic response, the diapause assay is challenging to perform in a high throughput manner. Therefore, we optimized the CCR assay as a phenotype for an organism’s ability to respond to the photoperiodic changes to perform the initial DGRP Genome-Wide Association Study (GWAS). We propose that our approach would identify genes involved in both photoperiodism and the circadian clock or only in photoperiodism. Our current study identified 32 candidate photoperiod genes from which we assessed the photoperiod response of the top 4 genes using CCR. We further characterized the role of allelic mutants of *PKCdelta* in photoperiodism using CCR, diapause, and molecular techniques.

## Results

### Variation in CCR in the DGRP

To test our hypothesis that genetic variations exist in photoperiodic response in *D. melanogaster*, we analyzed the CCR phenotype in DGRP, and found significant variations in CCR among DGRP strains (Figure 1, S1). The distribution of Cohen’s D values in DGRP, Shapiro-Wilks p = 3.763e-09, and W = 0.9194 (Figure 1A). To determine the differential response of each DGRP genotype to the two different photoperiods, CD values were calculated for each genotype (Materials and Methods). The majority of the DGRP strains fall under medium effect size (n = 91), large effect size (n = 61), and small effect size (n= 48) (Figure 1B). There is more variation in CD values among the large effect size DGRP strains than the strains that fall under medium and small effect size. The overall average CCR in LD is 941 sec while CCR in SD is 1095 sec in the whole DGRP population. Two representative strains are shown along with their recovery time under SD and LD (Figure 1C). DGRP109 has a delayed recovery time under SD in comparison to that of LD, while DGRP189 has a faster recovery time in SD compared to LD. To determine how much of the variation is due to genetic factors in the DGRP population, we calculated the heritability. Heritability (*h*^2^) for the long day condition was 0.51 and *h*^2^ for the short-day condition was 0.45. It was previously reported that there is a sex difference in the CCR of *Drosophila* (13). However, there was no strong correlation between sex and CCR in our experimental condition. The data from both sexes were grouped together for the GWAS analysis (Figure S2).

**Figure 1.**
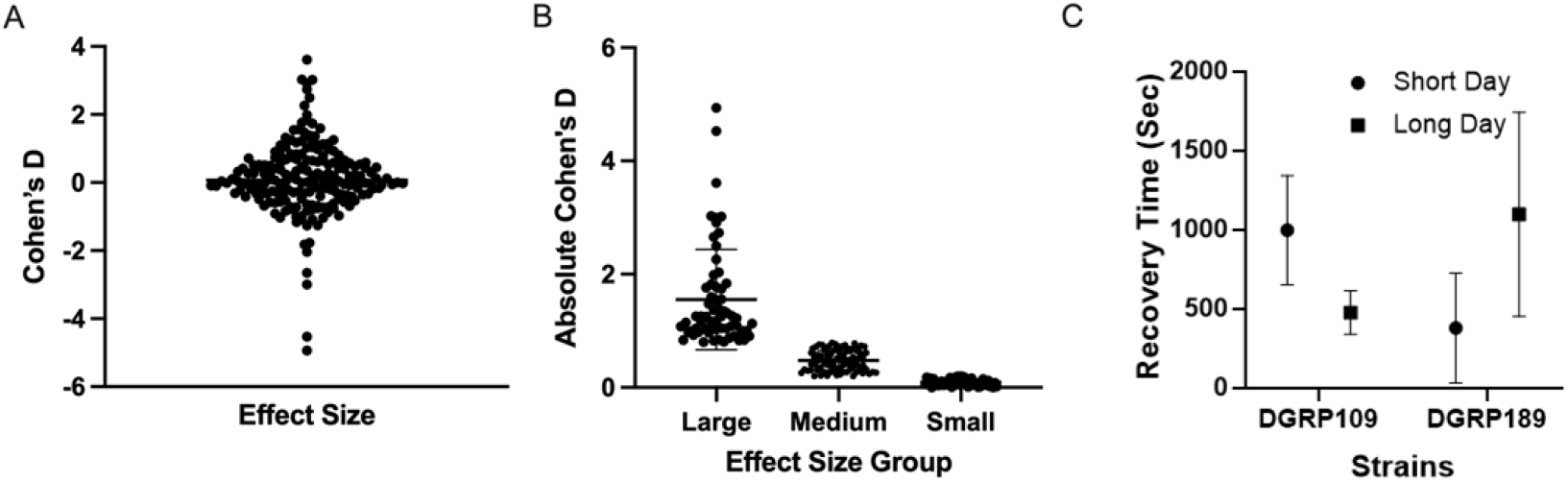
Variation in chill coma recovery in the DGRP population. A. The overall distribution of the CD values within the DGRP population, Shapiro-Wilks normality test p = 3.763e-09, p < 0.05, and W = 0.9194. B. The Cohen’s D (CD) of the DGRP population, CD = (CCR_SD_ - CCR_LD_)/average (STDV_CCR_SD_, STDV_CCR_LD_). CD > 0.8 is large effect size, CD of 0.5 is medium effect size, and CD < 0.2 is small effect size. Thick horizontal lines represent mean and thin lines represent standard deviation. C. The CCR of the two representative strains from the DGRP population, DGRP109 and DGRP189 under Short Day (SD) and Long Day (LD) photoperiods.

### Candidate Genes identified by GWAS

We characterized the top four candidate genes that are strongly associated with photoperiod CCR responses after narrowing down from 32 potential hits of GWAS analysis (Figure S3). These genes include, *Glucuronyltransferase-P* (*GlcAT-P*), *Brain-specific homeobox* (*Bsh*), *Protein Kinase C delta* (*Pkcdelta*), and *Diuretic hormone 31 Receptor* (*Dh31-R1*). *GlcAT-P* belongs to the glycosyltransferase family which are involved in catalyzing the conjugation of glucuronic acid from UDP-glucuronate to an acceptor. This particular gene is involved in the biosynthesis of L2/HNK-1 carbohydrate epitopes on both glycolipids and glycoproteins (24). *Bsh* encodes a putative transcription factor, and it is needed for the specification of neural identities in the adult brain. The rat ortholog of *Bsh* (*Bsx*) is suggested to be under circadian regulations and is expressed in the mature pineal gland of rat (25). *Dh31-R1* is *Diuretic hormone 31 Receptor* which encodes a protein that exhibits diuretic hormone receptor activity, and it is involved in G protein-coupled receptor signaling pathways. *Dh31-R1* is also known to be involved in time measurement and activity rhythms through the circadian clock system (26). *PKCdelta* belongs to a family PKC consisting of serine/threonine kinases which are activated by extracellular signaling. PKCs are divided into subgroups of Classical PKCs, Novel PKCs, and Atypical PKCs. *PKCdelta* belongs to the class of Novel PKCs which happen to be known for their two C1 domain and activation via 1,2-diacylglycerol (DAG) and phosphatidyl-serine (PS). While the cPKC family requires DAG and PS and aPKC only requires PS as a cofactor for activation (27,28). We identified two independent SNP variants in *PKCdelta*, which strongly suggest that the alleles of *PKCdelta* is associated with the variation of CCR in the DGRP population.

### Photoperiodic responses of the knockouts of the candidate genes

To further characterize the candidate genes, first, we analyzed CCR response of the knockout strains; *PKCdelta* mutant PKC*δ*^*e*04408^ (BDSC# 18258), *GlcAT-P* mutant *GLCAT* – *P*^0027–*G*4^ (BDSC#62582), *Bsh* mutant *Bsh*^*G*3075^ (BDSC#27467) and *Dh31-R1* mutant *Dh*31 – *R*^*MB*00175^ (BDSC#22718). As a control, one DGRP parent strain DGRP208 (WT) was used to compare between the Cohen’s D values. For this analysis equinox (EQ) condition, a 12:12 hr light:dark cycle was used along with SD and LD photoperiods. The CD was obtained by subtracting (or normalizing) the EQ photoperiod CCR from SD & LD CCR. The WT strain has a negative CD value of −0.69 (Figure 2A). The *Dh31-R1* mutant has a similar photoperiod response to the WT with negative CD values in both SD and LD photoperiod. *Bsh* mutant also has negative CD value though the effect is not as exaggerated. Both *GlcAT-P* and *PKCdelta*;Pbac mutants has positive CD values (Figure 2A). To determine how different these mutants’ photoperiod responses were from the WT control, the CD value of the WT was subtracted from the CD values of the mutant, the final value is termed Genetic Effect Size (GES) (Figure 2B, Materials and Methods). *Bsh*, *GlcAT-P* and *PKCdelta*:Pbac have different photoperiod responses compared to the WT with *PKCdelta* being the most different. The GES of *PKCdelta*:Pbac is not significantly different between LD and SD. However, *GlcAT-P, Bsh*, and *Dh31-R1* have a difference in GES between LD and SD (Figure 2B).

**Figure 2.**
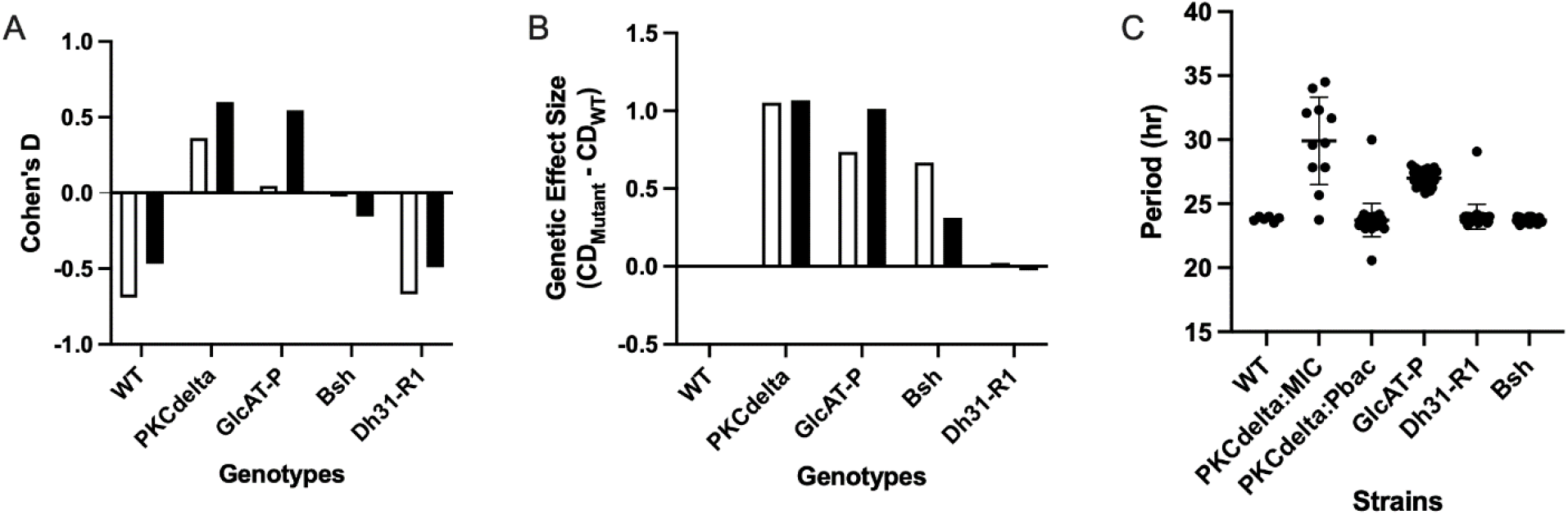
The photoperiod response and clock period phenotype of the top four candidate knockout strains A. Cohen’s D values of the knockout mutants of the four candidate genes under LD and SD photoperiods compared to the wild type DGRP 208 strain (WT). The CD is normalized to equinox (EQ) photoperiod (Materials and Methods). The WT, *Dh31-R1* and *Bsh* have negative CD in LD (open) and SD (filled), while *PKCdelta* mutant *PKCdelta*:Pbac and *GlcAT-P* have positive CD under both conditions. B. The Genetic Effect Size of the candidate gene mutants, *PKCdelta*:Pbac and *GlcAT-P* mutants show a different photoperiod response compared to the WT under LD (open) and SD (filled). C. The circadian period of the candidate gene mutants compare to the WT with the period of 24 hr. *GlcAT-P* has a significantly longer period compared to WT (t-test: p < 0.05, p = 1.16E-13) and *PKCdelta*:MIC mutant of the *PKCdelta* has longer period compared to the WT (t-test: p < 0.05, p = 1.28E-3). Error bars represents standard deviation. N > 20.

Next, we tested the hypothesis that there are shared genetic regulators between the circadian clock and photoperiodism. We examined the period of the candidate photoperiod knock out genes using WT as a control (Figure 2C). Compared to the WT which had a period of 24 hr, *GlcAT-P* and *PKCdelta*:MIC mutants have a significantly longer period (t-test: p < 0.001, p<0.01). We concluded that *GlcAT-P* and *PKCdelta*:MIC genes play a role in both circadian clock regulation and photoperiodism. We decided to investigate *PKCdelta* further for three reasons. The two *PKCdelta* mutant alleles showed a significantly different photoperiodic CCR response from that of the WT (Figure 2B), one of the allelic mutants of *PKCdelta* had an altered circadian period (Figure 2C), and GWAS analysis showed that there are two independent SNP variants associated with the photoperiodic CCR response for this gene (Figure S3 and File S3).

### PKCdelta Candidate Gene

*PKCdelta* is a kinase known to be involved in cellular signaling pathways like cell growth, apoptosis, tumor promotion and carcinogenesis (28). We wanted to identify if there are SNPs that could lead to strains having a large Positive CD and large negative CD. So, we examined two parental DGRP strains one with a large positive CD value (DGRP379) and another with a large negative CD value (DGRP391). *PKCdelta* transcribes three transcripts, RC, RD, and RE (Figure S6). The transcripts RC and RD are similar to each other except with a slight difference in the UTR region; the RD isoform has a shorter UTR region than the RC isoform. The RC and RD transcript have regulatory domain which contain C2, C1 and pseudosubstrate domains toward the N-terminal. Toward the C-terminal there is a catalytic kinase domain which is shared with all three isoforms (Figure S6). There are two different transcription start sites, one that starts the beginning of isoforms RD and RC, and the other starts at the RE isoform. The RE isoform is a nested gene with its transcription start site residing in one of the introns of RC and RD transcripts (29).

We found that there are two non-synonymous SNPs in the exon 6 which resides in the RE transcript in the DGRP379 strain but not in DGRP391. The first SNP position is at 12445745: Guanine to Adenosine, which changes Alanine to Threonine. The second SNP position is at 12446453: Guanine to Cytosine, which changes Alanine to Proline. The first SNP position is in a ployQ region, which is conserved in the *Drosophila* species. The second SNP causes a repeat of the sequence surrounding the region (Figure S7A, S6B). We found that RE isoform is a nested gene which is only conserved in the *Drosophila* genus (Figure S8).

### Diapause and CCR in PKCdelta mutant alleles

To further explore the role of *PKCdelta* in photoperiodism, we performed the diapause assay using the *PKCdelta*:Pbac and *PKCdelta*:MIC allelic mutants under different photoperiodic conditions. The *PKCdelta*:Pbac mutant has an insertion of a *piggy bac* element in an intron close to the 3’ end of the gene. The *PKCdelta*:MIC mutant has an insertion in the 5’ end of the gene and contains a minos-based construct which was created by the Gene Disruption Project (GDP) (30) (Table S4). We found that both mutant alleles have a mutation in the catalytic domain of the enzyme *PKCdelta* (Figure S10, Table S3 and S4)

Using WT as a control, the diapause assay was carried out under SD and LD conditions (Figure 3A). As expected, the WT has increased diapause incidences under SD conditions compared to the LD condition (Z-test, p < 0.05). There was a significant difference in *PKCdelta*:Pbac mutant’s diapause response compared to that of WT in both photoperiods (Z-test, p = 0.0014 (LD), p = 0.01 (SD)). The *PKCdelta*:MIC mutant’s diapause response compared to WT in each photoperiod is not significantly different (Z-test, p = 0.27 (LD), p = 0.30 (SD)). However, there is a significant difference in *PKCdelta*:Pbac LD and SD diapause response (Z-test, p = 0.002), and in WT (Z-test, p = 0.007), but not in *PKCdelta*:MIC (Z-test, p = 0.29) (Figure 3A). The diapause response in *PKCdelta*:MIC mutant is not statistically significant from that of the WT. We interpreted this as the mutant allele *PKCdelta*:MIC cannot distinguish SD from LD because the diapause response between photoperiods is not significantly different (Z-test: p = 0.29).

**Figure 3.**
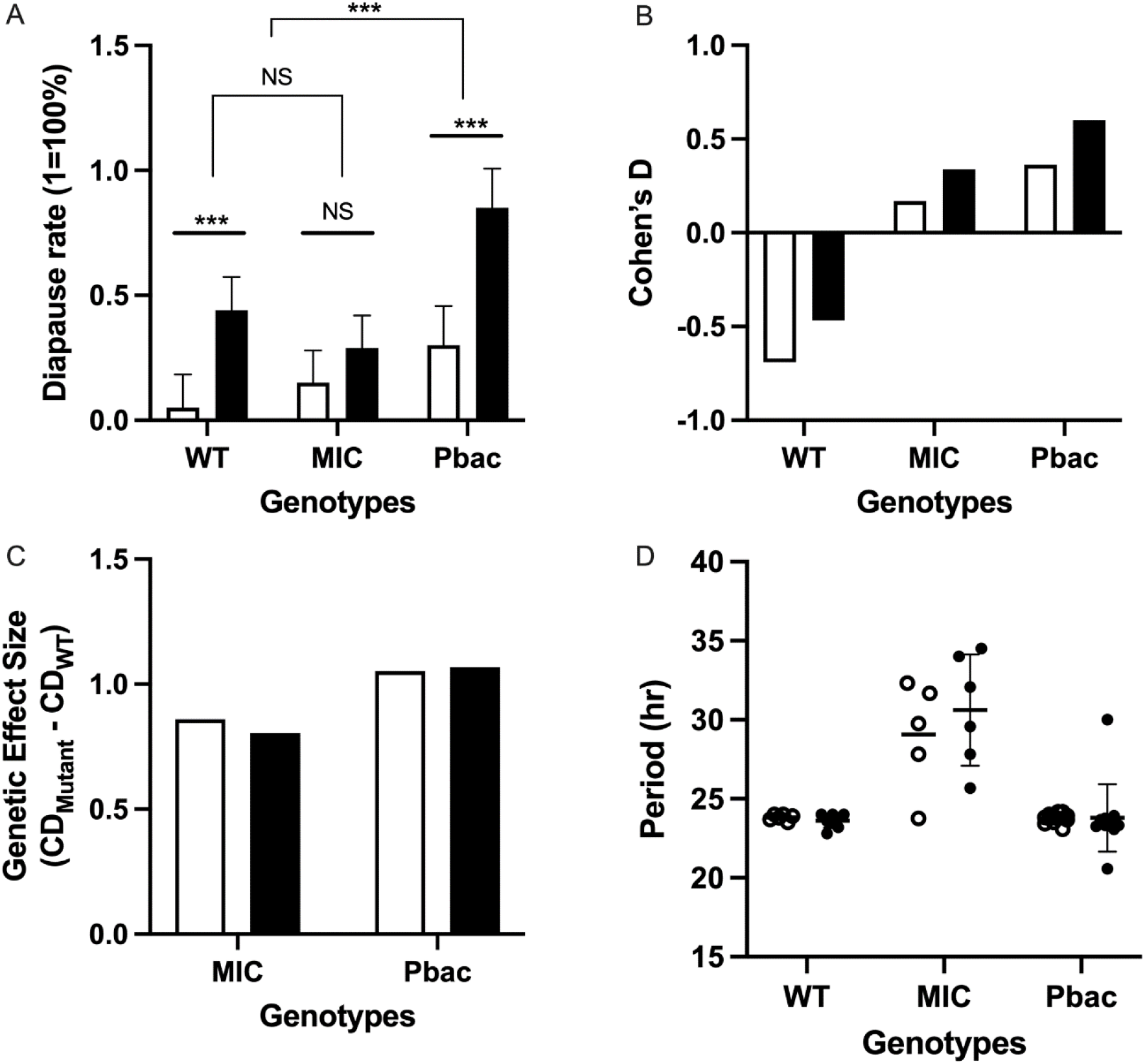
The photoperiod and period response of *PKCdelta*:MIC and *PKCdelta*:Pbac mutants compared to WT. A. The diapause response of candidate gene *PKCdelta* mutants compared to WT in LD (open) vs SD (filled). There is a significant difference in *PKCdelta*:Pbac mutant diapause response, compared to WT in both photoperiods (Z-test: p= 0.0014 (LD), 0.010 (SD)). *PKCdelta*:MIC diapause response compared to WT in each photoperiod is not significantly different (Z-test: p = 0.27 (LD), 0.30 (SD)). However, there is a significant difference in *PKCdelta*:Pbac LD and SD diapause response (Z-test p = 0.002) and in WT (Z-test: p = 0.007), but not in *PKCdelta*:MIC (Z-test: p = 0.29); n=20-25 flies. B. Cohen’s D of *PKCdelta*;Pbac and *PKCdelta*;MIC compared to WT, the CD values were obtained by subtracting EQ CCR response from LD (open) and SD (filled) photoperiod response. WT has a negative CD, while *PKCdelta*:MIC and *PKCdelta*:Pbac are positive. C. GES of the mutants compared to the WT, both mutant alleles show significantly different photoperiod response from that of the WT, LD (open), SD (filled). D. Circadian period of the *PKCdelta* mutants compared to that of WT separated by homozygous females (open) and hemizygous males (filled). Error bars represent SD, 10 < n < 20, homozygous vs hemizygous (t-test, p > 0.05).

To determine the necessity of *PKCdelta* for the photoperiodic CCR response, we preformed the CCR assay using the allelic mutants of *PKCdelta* (Figure 3B). The CD values of *PKCdelta*:Pbac and *PKCdelta*:MIC are positive compared to WT. However, *PKCdelta*:MIC have lower CD values from that of *PKCdelta*:Pbac. The GES of the mutants compared to the WT show that *PKCdelta*:Pbac have significantly different photoperiod response (Figure 3C).

Since *PKCdelta* is located on the X-chromosome, both of the *PKCdelta* mutants contain hemizygous males and homozygous females. The CCR was performed to determine whether there will be a difference in photoperiod response between hemizygous and homozygous flies under SD and LD photoperiod. Both *PKCdelta*:MIC and *PKCdelta*:Pbac mutants show a difference in photoperiod response between hemizygous and homozygous flies (Figure S9). To determine if there was a difference in period phenotype between hemizygous and homozygous flies in the *PKCdelta*:Pbac and MIC mutants, we performed the period assay. We found no significant difference between the period phenotype between hemizygous and homozygous flies (Figure 3D). WT flies have an expected period of 24 hr with no difference between hemizygous and homozygous flies (t-test, p > 0.05). *PKCdelta*:MIC hemizygous flies have a period of 30 hr and homozygous flies have a period of 29 hr but there was no statistically significant difference between hemizygous and homozygous flies (t-test, p > 0.05). There was no significant difference between hemizygous and homozygous flies of *PKCdelta*:Pbac in the circadian period (t-test, p > 0.05). Based on these data together, we concluded that MIC mutant allele affected the ability to distinguish between LD from SD, and also altered the circadian period.

Next, we predicted that *PKCdelta*:Pbac has knockdown of all three transcripts of *PKCdelta*, while *PKCdelta*:MIC has knockdown of RC & RD based on the reported locations of the insertion. The RE isoform is believed to be mainly responsible for photoperiod response because the *PKCdelta*:MIC mutant shows a decrease in the photoperiod response in the CCR and diapause assays. We also predicted that the transcripts RC and RD could play a role in interpreting photoperiod response, though they might not be major players in interpreting photoperiods. These predictions were then confirmed by performing a qPCR experiment using the *PKCdelta*:MIC and *PKCdelta*:Pbac mutants under SD and LD condition (Figure 4).

**Figure 4.**
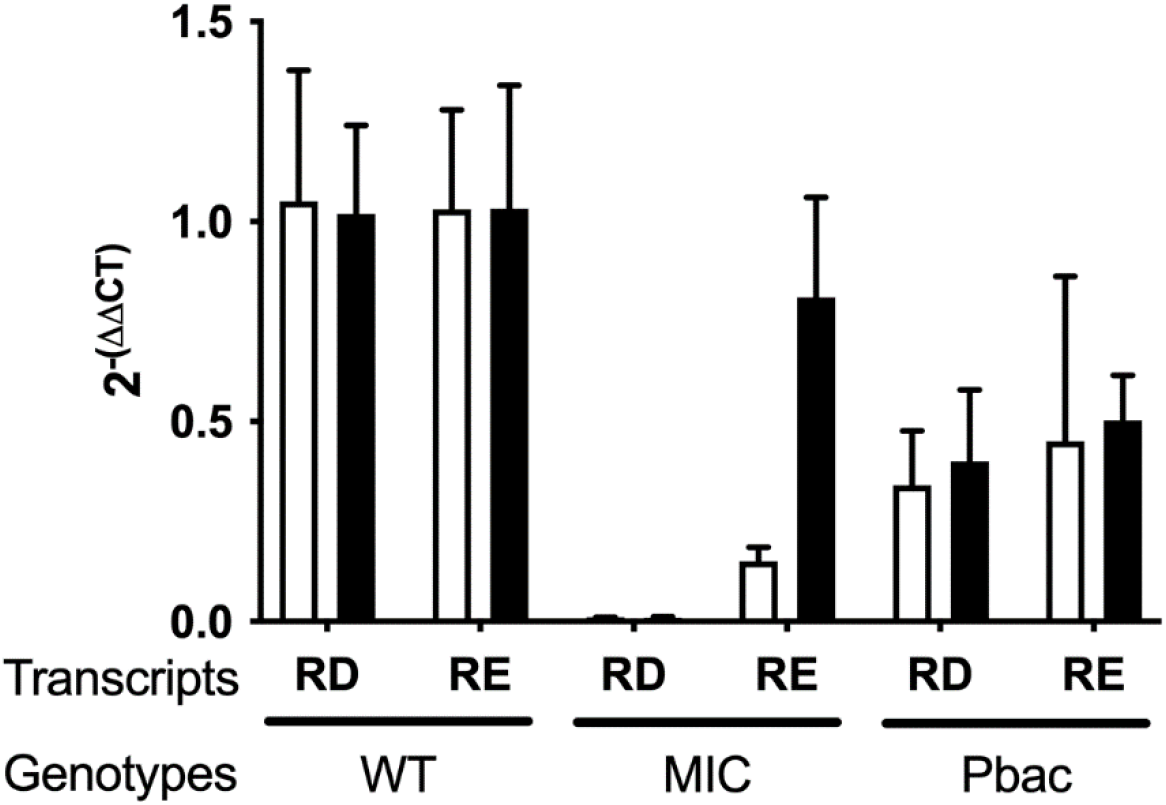
Expression level of *PKCdelta* allelic mutants under LD (open) and SD (filled) compared to DGRP 208 WT. In the WT flies there is no difference in expression of RD and RE transcripts under LD and SD (n=3-5, t-test, p > 0.05). In *PKCdelta*:MIC the expression of RD is significantly reduced compared to WT (n=3-5, t-test, p<0.001). However, there is a reduction of RE transcript under LD compared to SD in *PKCdelta*:MIC (n=3-5, t-test, p < 0.001). In the *PKCdelta*:Pbac background, the expression of RD and RE is overall reduced compared to WT (n=3-5, t-test, p<0.05). There was no statistical difference in the expression levels under LD and SD photoperiods (n=3-5, t-test, p > 0.05).

### Expression of PKCdelta

We measured the expression levels of *PKCdelta* under different photoperiod conditions (Materials and Methods). WT flies under LD and SD photoperiods did not show statistically significant change in *PKCdelta* RD and RE transcript levels (t-test, p > 0.05). *PKCdelta*:MIC which was predicted to have reduced expression of RD and normal expression of RE, show overall decrease in expression of RD in both LD and SD (t-test, p > 0.05). However, there is an alteration in the expression levels of the RE between the photoperiod conditions in *PKCdelta*:MIC. The *PKCdelta:*MIC mutant under LD show a significant decrease in its expression level from that of SD (t-test, p < 0.001). *PKCdelta*:Pac mutant show an overall decrease in RD and RE expression from that of WT (t-test p < 0.05). Though there is no significant difference in the expression levels of RD and RE between the photoperiods in *PKCdelta*:Pac mutant, (t-test p > 0.05). This suggests that there is an alteration in the expression of the RE transcript in the *PKCdelta*:MIC mutant due to long day photoperiod condition. Based on this data and the diapause data, we concluded that RD transcript could be involved in distinguishing between SD and LD. Since *PKCdelta*:MIC which has reduced expression of RD transcript show no difference in the diapause response between photoperiods (Figure 3A & 4). Alternatively, the *PKCdelta*:Pbac mutant which has reduced expression of both transcripts shows a greater proportion of diapause response compared to WT in both LD and SD suggesting that expression *PKCdelta* and diapause response is negatively correlated in this mutant.

Next, we reasoned that the GWAS analysis of the DGRP CCR response itself as a trait under LD and SD conditions will lead us identifying the downstream genes involved in CCR. We performed the GWAS analysis on the CCR trait as a phenotype rather than the photoperiodic response (CD values) as a trait discussed earlier (Files S4 - S7). There were six genes that were highly associated with the CCR phenotypes both in SD and LD conditions: *Sif, Comm2, Bru-3, Fife*, *CG32111* and *Spab*. *Sif* is involved in actin cytoskeleton organization; visual perception and it localizes to the synapse. *Comm2* is essential for nerve cord development and axon guidance. *Bru*-3 is known to mediate mRNA translation repression. While *Fife* is responsible for encoding synaptic scaffolding protein, which promotes neurotransmitter release. *CG32111* is a long non-coding RNA though much information is not known about this gene. *Spab* is a protein coding gene from the neuropeptide gene group, however not much is known about this gene (Flybase.org). As expected, most of the genes responsible for the CCR response play a role in the nervous system. However, *Sif* and *CG32111* are known to be a target of the circadian clock transcription complex CLK/CYC, suggesting that CCR photoperiod response could be regulated by the clock (31). *Sif* is also known to interact with *Fas2* which than interacts with *PKCdelta* (32).

## Discussion

In this study, we were able to establish that there is a variation in photoperiod response measured by the CCR phenotype within the DGRP population. By calculating the effect size (Cohen’s D value) for each DGRP genotype we determined how each genotype’s photoperiod response was different from one another. The GWAS analysis on CD values allowed us to narrow down the 32 photoperiod candidate genes. We chose 4 top candidate photoperiod genes for genetic analysis. The allelic mutant of *PKCdelta* showed a significantly different photoperiod response from WT. To test the necessity of the gene for the photoperiodic response, we analyzed two different mutant alleles: *PKCdelta*:Pbac and *PKCdelta*:MIC. By performing a qPCR assay under LD and SD it was confirmed that *PKCdelta*:Pbac has decreased expression of RD and RE transcripts of the gene. The *PKCdelta*:MIC mutant which show reduced expression of the RD transcript, show a photoperiod effect under LD in the RE transcript levels. The CCR assay showed that *PKCdelta*:Pbac mutants have a greater effect size compared to *PKCdelta*:MIC. Finally, to test the hypothesis whether there is an overlap in the genetic mechanisms between the circadian clock and photoperiodism, the circadian periods of our candidate genes were analyzed. *GlcAT-P* knockout and *PKCdelta*;MIC mutants had a significantly longer period phenotype from the WT. Our data supports the view that *PKCdelta* is involved in both circadian clock and photoperiodism.

Diapause and CCR are two different output pathways that are under the photoperiod regulation in insects. Our data showed that the two different mutant alleles of *PKCdelta* manifested the altered photoperiod response in different ways. The flies with *PKCdelta*:MIC allele could not distinguish LD from SD, whereas the flies with *PKCdelta*:Pbac allele had an elevated instances of diapause under both photoperiods (Figure 3A). The CCR data suggests that the *PKCdelta*:Pbac mutation plays a role in increasing the sensitivity to photoperiod in the flies overall. Both the mutants have positive CD values indicting their recovery time was faster than that of WT. While *PKCdelta*:Pbac compared to *PKCdelta*:MIC has a greater effect size (Figure 3B).

Taken together, we have a working model for the photoperiodism in *D. melanogaster*. The photoperiod candidate gene *PKCdelta* is hypothesized to be regulated by the clock. There are certain genetic components that are under clock control which interact with *PKCdelta*. The circadian transcription complex CLK/CYC is known to directly regulates the *Mef2* transcription which then has been identified to target *Fas2*. *Mef2* is known for its role in myogenesis and development (33). However, *Mef2* is shown to be present in clock neurons and fluctuation in *Mef2* are rhythmic (33). *Mef2* is a major component of the CLK/CYC, which is important for circadian behavior of neuronal morphology. A Chip-Chip analysis determined that *Fascilin2* (*Fas2*) is a direct target of *Mef2* (33). *Fas2* is known to be involved in neural cell adhesion, and there is a rhythmic oscillation of *Fas2* in pigment-dispersing factor (PDF) cells types, suggesting that *Fas2* is under circadian clock control (33). *PKCdelta* is proposed to interact with *Fas2* (32). PDF is known to effect *Drosophila* circadian clock where certain mutations in PDF can shorten the period of certain clock neurons while lengthening the period of other neurons (34). The diapause assay showed that *PKCdelta*:MIC mutants lose the ability to distinguish between day-lengths. *PKCdelta*:MIC mutant’s loss of photoperiod sensitivity is similar to the well-studied *PDF*^01^ mutant (35). Our data reject the model that circadian clock and photoperiodism are separate entities with no shared regulations. The current study sheds light onto potential genes that could be further studied to characterize the mechanisms of photoperiodism, and its relationship to the circadian clock.

## Materials and Methods

### Fly stocks

We purchased the DGRP strains from the Bloomington Drosophila Stock Center (Indiana University, Bloomington, IN). In the laboratory, the fly stock was maintained at room temperature (25 □) and in 12L:12D photoperiod conditions. The following are the *PKCdelta* mutants BDSC#18258 Pbac[RB]PKCdelta[e04408], BDSC#59433 Mi[MIC]PKCdelta[MI13377]. The BDSC#62582 is *GlcAT-P* knockout mutant Pbac[GlcAT-P]0027-G4, BDSC#27467 Bsh mutant EPbsh[G3075]. BDSC#22718 is the *Dh31-R1* knockout mutant ET1Dh31-R1[MB00175].

### Photoperiod conditions and Chill coma recovery

Chill Coma Recovery (CCR) is a traditional method used to assess the photoperiod phenotype in *Drosophila* species. CCR is expressed as the time (in seconds) it takes an organism to recover from a paralysis like state after being kept under extremely low temperature. *D. melanogaster* enter the chill coma at 7.0 ± 0.9 °C (36). It has been clearly demonstrated that flies raised under short-day (SD) photoperiod conditions compared to long-day (LD) photoperiod conditions have an advanced (or shorter) recovery time (37). DGRP fly lines were reared under LD and SD in a chamber, 25 ± 0.5 □ and 70% humidity. The LD consisted of 16 hours of light and 8 hours of dark (16h:8h), while the SD consisted of 8 hours of light and 16 hours of dark (8h:16h). The flies were reared under these conditions for two weeks. Chill coma was then performed by placing the flies in 4 □ for 60 minutes. Next, the flies were placed into 5 mm glass tubes and into the *Drosophila* activity monitors (DAM2, Trikinetics) to measure the time it takes for the flies to recover. For an accurate measure of the recovery time, the flies were placed in the middle of the glass tube near the IR sensor to capture the exact time the fly recovered from the coma. 200 DGRP lines were used for this assay which included 38 core parent strains and 162 homozygous offspring strains, 20 replicate per DGRP strain were used.

### Diapause Assay

The diapause assay was performed by surgically removing the ovaries after rearing female flies under different photoperiods. Previous research suggested that female *Drosophila* placed under SD enter reproductive diapause which results in underdeveloped ovaries. The SD chamber condition mimics the winter-like days when most organisms cease to reproduce to conserve energy, therefore, diapause is induced in female flies (9). Female *Drosophila* placed under LD condition do not enter diapause. The *PKCdelta* mutants, *PKCdelta*:Pbac and *PKCdelta*:MIC were reared under SD and LD photoperiodic conditions along with DGRP208 (WT) strain as a control. Newly hatched female flies were placed under the photoperiod conditions for 12 days, 25° C. Ovaries were dissected in 1x PBS, the image was taken with a light microscope (LEICA microsystems, LEICA EC3, Model:MEB126) under 4x magnification. Flies were considered to be in diapause when eggs in both the ovaries were under stage 8 of development.

### qPCR

The expression level of the candidate gene *PKCdelta* was characterized under SD and LD. Whole fly RNA extraction using the Zymo Research Direct-Zol RNA extraction kit (Cat#R2050) was prepared. The cDNA was prepared using the New England Biolabs (NEB) protoscript II first strand cDNA synthesis kit (Cat#E65605). For qPCR the Applied Biosytems PowerUp SYBER Green master mix was used (Cat#A25742) in the Applied Biosystems QuantStudio 6 Rlex Real-Time PCR system according to the manufacture’s manual. The following primer set was used; ACTIN served as a control, (Fw; GCAAGTACTCTGTCTGGATC) (Rv; CCAGCAGAATCAAGACCATCC). HK-5 primer set which targeted the RD transcript of *PKCdelta* (Fw; CGATCGATTGGGGTTTGCTGG) (Rv:GTGATGTGAGGATTTGTGTAGG), expected product size 283 bp. HK6 primer set targeted the RE transcript of *PKCdelta* (Fw; TCGAGCTGAAGACAACACCT) (Rv: CTTTCTTAGCTTCGACTTGGTT) expected to product size 160bp (Figure S5, Table S2).

### Cohen’s D

CCR has been used successfully by other groups as the phenotype reflecting photoperiodic response (13). Our primary interest is measuring the ‘differential’ response of each genotype to two different photoperiodic conditions. Cohen’s D (CD) is used to measure the effect size of the photoperiodic response in each genotype (38):

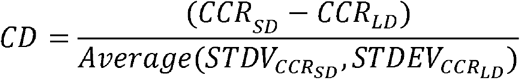

For each of the DGRP strains, the average CCR time was calculated from 20 replicates per strain in SD (CCR_SD_) and LD (CCR_LD_). The CD value of each genotype was calculated by subtracting CCR_LD_ from CCR_SD_, and divided by the average of the two standard deviations of CCR_LD_ and CCR_SD_. The CD values can be divided into three groups: large effect size, CD > 0.8; medium effect size, 0.8 > CD > 0.5; and small effect size, CD < 0.2. The CD values for the mutants were obtained by subtracting equinox photoperiod CCR value from that of SD or that of LD respectively. This was done to ensure that the data was normalized to the EQ photoperiod.

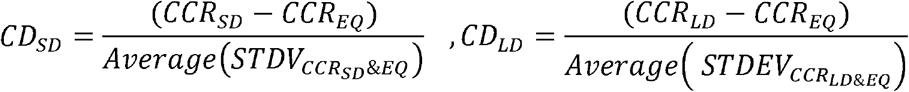

CD value provides directionality by positive or negative values. Thus, CD value shows if a photoperiodic condition provided a positive effect on the strain or a negative effect in the CCR assay.

### Genotype Effect Size

To measure the difference between the effect size of CD in different genotypes, we calculated the Genotype Effect Size (GES) by subtracting the CD value of the wild type from that of the mutant.

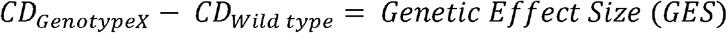

In calculating the GES values, DGRP208 (WT) was used as the baseline. GES value allowed us to determine which candidate gene had the most differential response to the photoperiod in comparison to that of the wild type.

### GWAS Analysis

After calculating CD for all the strains under different photoperiodic conditions, we performed the GWAS using the web-based tool provided by DGRP2 (21,22). The web-based tool requires inputting a phenotype file to run the GWAS analysis and the users can expect to retrieve all association results. The 1,881,979 select SNPs were analyzed for their association with the photoperiod response. All the variants from the GWAS were graphed in a manhattan plot with their chromosome on the X-axis and -log10(p) of the p values on the Y-axis. There are two significant thresholds -log10(1e-5), for “suggestive” association threshold and -log10(5e-8) for “genome-wide significance” threshold (Figure S3).

### Heritability

The HiLMM R package was used to calculate narrow-sense heritability of photoperiod response in the DGRP population. The HiLMM package provides an estimation of heritability with confidence intervals in linear mixed models.

## Data Availability

All DGRP lines are available at the Bloomington Drosophila Stock Center, and the genotype information of all lines are available at http://dgrp2.gnets.ncsu.edu. The raw CCR and CD data for all DGRP lines are available under File S1. The Files S2, S4, & S6 contain the top GWAS associations along with qq-plots, and input data used to obtain the GWAS results from CD, LD, and SD data sets. The Files S3, S5, & S7 contain all associations found in the GWAS study using the CD, LD and SD data sets. All Supplemental Materials are available at figShare portal, gsajournals.figshare.com.

## Acknowledgements

The authors are grateful for Drs. Nir Yakoby, Nathan Fried, and Charot Rodeget for critical reading and insightful suggestions. We are grateful for the Lee lab members, especially Sienna Casciato for helpful discussions. The authors would also like to thank Cody Stevens for helping us navigate through some of the challenging aspects of the project. The work was funded by the NIH MARC U-STAR, T34GM127154.

## Supplemental Material

**Figure S1.** The distribution of core 38 set DGRP parent strains.

**Figure S2.** Sex difference in CCR assay among the select DGRP strains.

**Figure S3.** Manhattan plot for the GWAS analysis of DGRP.

**Figure S4.** The Gene enrichment results of the 32 hits using flymodEnrichr.

**Figure S5.** Diagram of the two *PKCdelta* mutant alleles, Pbac and MIC.

**Figure S6.** Overview of the *PKCdelta* gene structure and the position of the oligos for PCR products.

**Figure S7.** Sequence variations in *PKCdelta* RE transcript.

**Figure S8.** Phylogenetic analysis of *PKCdelta* RE transcript.

**Figure S9.** Cohen’s D and Genome Effect Size of *PKCdelta* mutants.

**Figure S10.** The regulatory and catalytic domains of *PKCdelta*.

**Table S1.** The top 32 hits of the GWAS analysis sorted by the significant p values.

**Table S2.** Oligos used in the current study.

**Table S3.** Predicted changes *PKCdelta*:Pbac mutant.

**Table S4.** Predicted changes in *PKCdelta*:MIC mutant.

**File S1**. Raw CCR and Cohen’s D data for all the DGRP lines. It’s divided into three tabs; Cohen’s values, CCR in LD and CCR in SD.

**File S2**. TOP GWAS hits along with images of QQ plots.

**File S3**. SNP associations found in the GWAS study.

**File S4**. TOP GWAS hits using the CCR data in LD.

**File S5**. All GWAS hits using the CCR data in LD.

**File S6**. TOP GWAS hits using the CCR data in SD.

**File S7**. All GWAS hits using the CCR data in SD.

