## Supplementary figures and images for "Roles of *PKCdelta* in Photoperiod and Circadian Regulations in *Drosophila melanogaster*"

### Figure S1

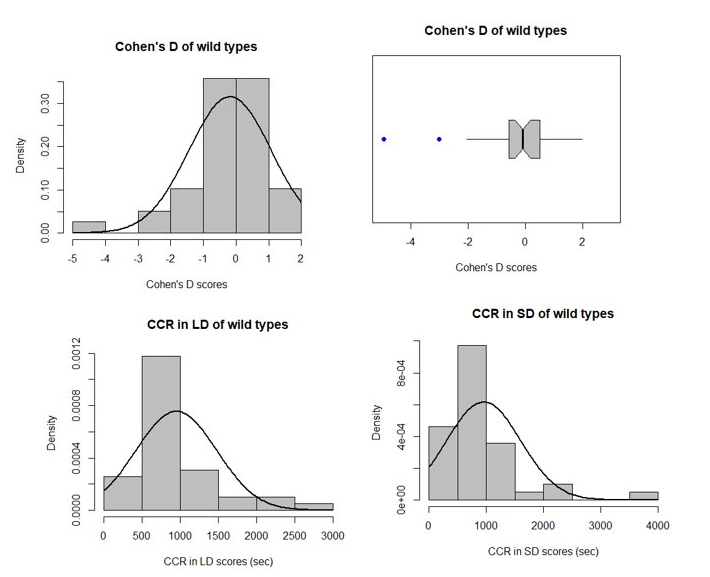

**Figure S1.** The distribution of core 38 set DGRP parent strains.

### Figure S4

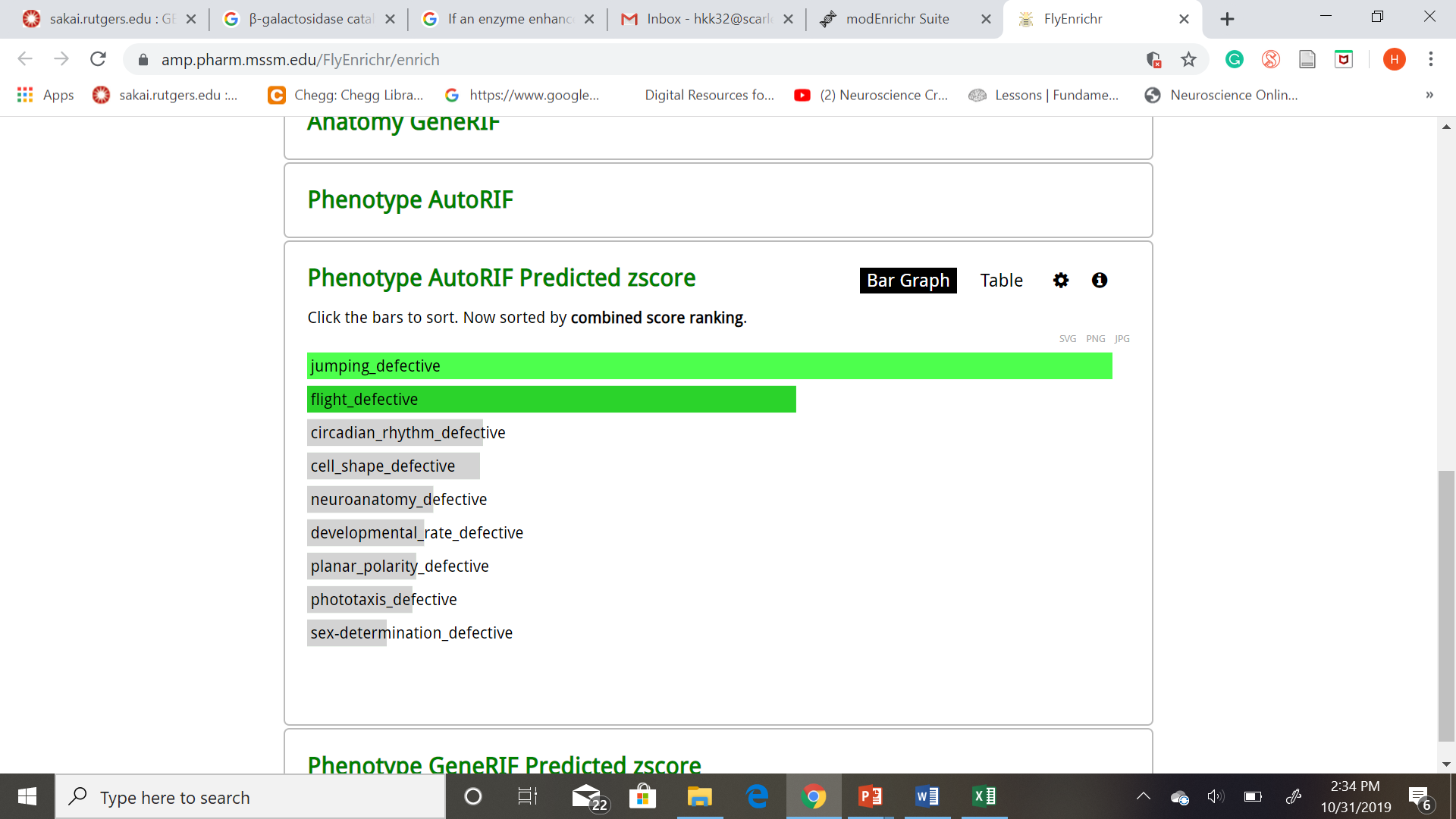

Enriched genes

**Figure S4.** The Gene enrichment results of the 32 hits using flymodEnrichr.
