## Supplementary material for "Roles of *PKCdelta* in Photoperiod and Circadian Regulations in *Drosophila melanogaster*": Figure S2

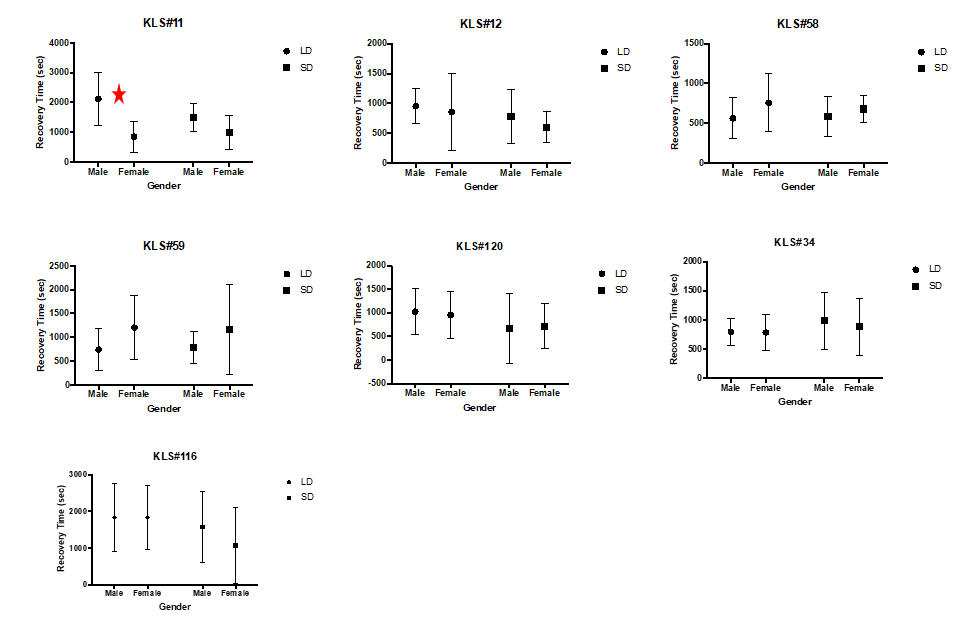

**Figure S2.** Sex difference in CCR assay among the select DGRP strains.

To determine if this sex difference would be prevalent in the DGRP population, 7 different genotypes, 20 males and 20 females under two different photoperiod conditions were assessed for their CCR response. By conducting pairwise analyses between the male and female flies it was found that only one genotype under LD showed statistical significance. Out of 14 strains tested, only one showed difference in recovery time KLS#11 under SD, overall, there was no significant difference the CCR between female and male DGRP flies.
