## Supplementary material for "Roles of *PKCdelta* in Photoperiod and Circadian Regulations in *Drosophila melanogaster*": Figure S3

*GWAS Analysis for Photoperiodism*

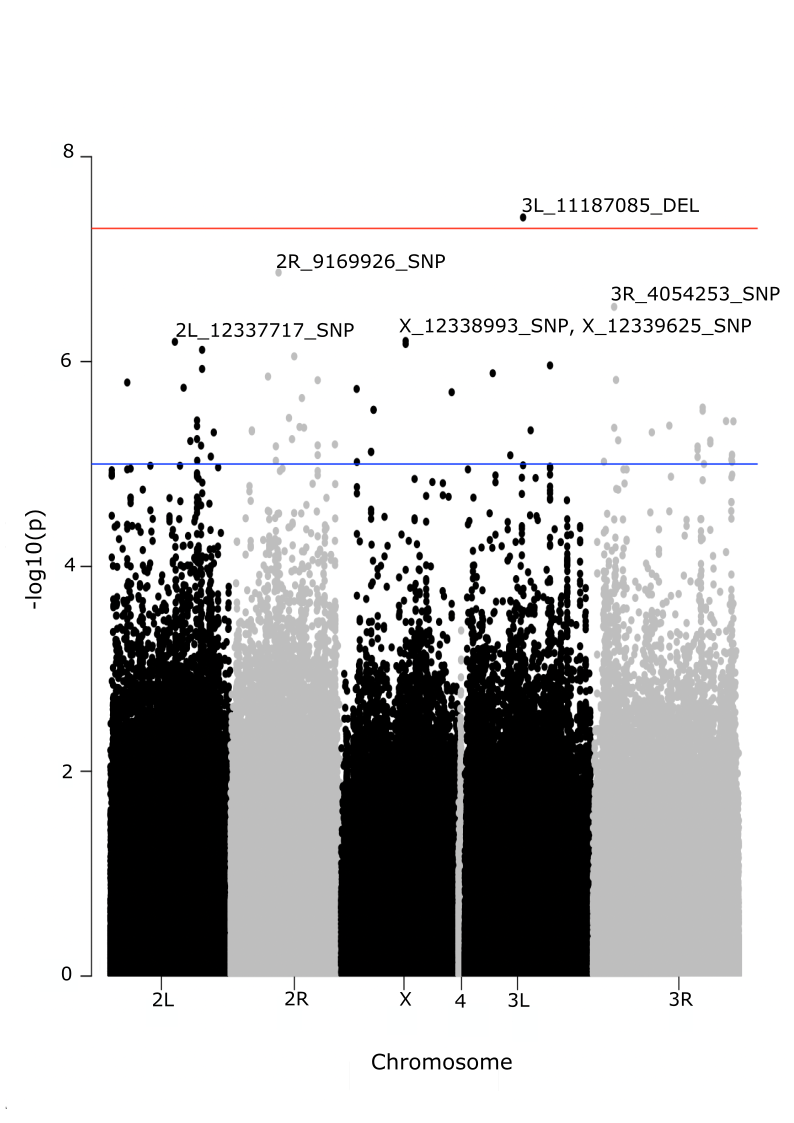

Figure S3. Manhattan plot for SNPs in the GWAS analysis. Each point represents a single SNP, and the height of the SNP represent the strength of association with photoperiod (CCR) response expressed as –log10(p). The horizontal lines represent the genome-wide Bonferroni significance threshold (blue line p = -log10(1e-5), red line p = -log10(5e-8). The variant 3L_11187085_DEL is associated with GlcAT-P, 2R_9169926_SNP is associated with Dh31-R1, X_12338993_SNP and X_12339625_SNP are associated with PKCdelta. The variants 2L_12337717_SNP and 3R_4054253_SNP are not associated with any particular gene since they are in the intergenic regions. We also identified the 32 most promising variants strongly associated with the photoperiod response (Table S1). The enrichment analysis of these candidate genes shows a list of biological phenotypes including jumping, flight phenotypes, and circadian rhythms (Figure S4).
