## Supplementary material for "Roles of *PKCdelta* in Photoperiod and Circadian Regulations in *Drosophila melanogaster*": Figure S5

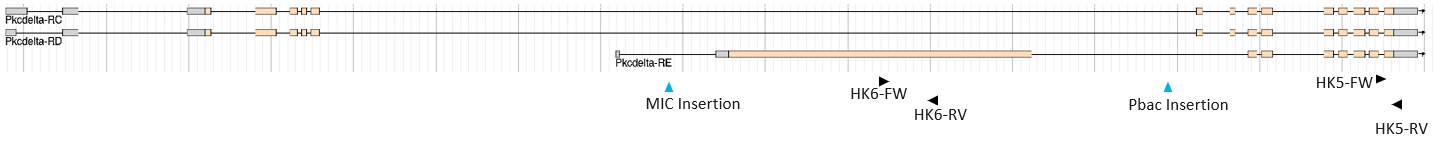

**Figure S5.** Diagram of the two *PKCdelta* mutant alleles, Pbac and MIC.

The primers used for qPCR assay to determine the expression level of *PKCdelta* allelic mutants under SD and LD are shown.
