## Supplementary material for "Roles of *PKCdelta* in Photoperiod and Circadian Regulations in *Drosophila melanogaster*": Figure S6

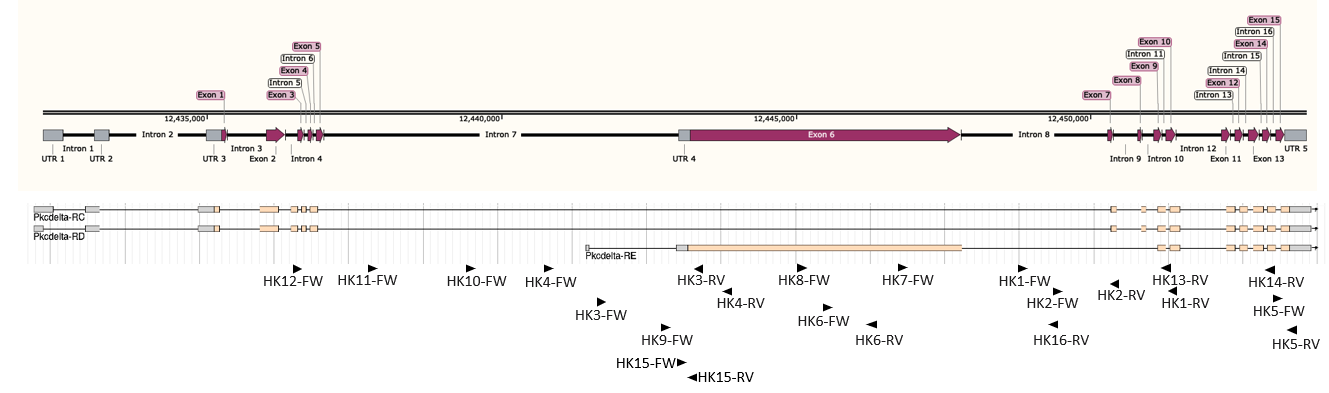

**Figure S6.** Overview of the *PKCdelta* gene structure and the position of the oligos for PCR products. The drawing modified from flybase.com.
