## Supplementary material for "Roles of *PKCdelta* in Photoperiod and Circadian Regulations in *Drosophila melanogaster*": Figure S7

A

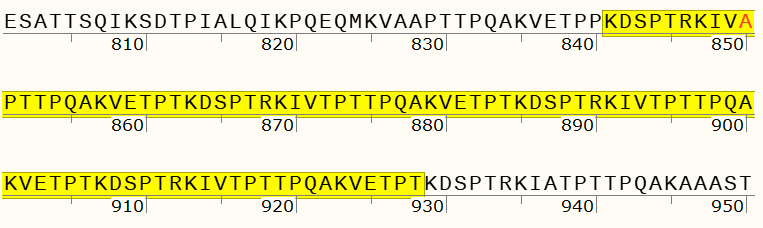

B

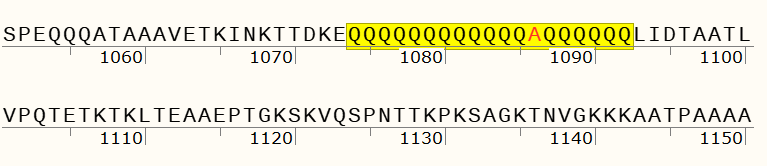

**Figure S7.** Sequence variations in *PKCdelta* RE transcript.

The location of the nonsynonymous mutations in exon 6 *PKCdelta* RE transcript. A. The location of SNP in DGRP379 is at 12445745: Guanine to Adenosine, which changes Alanine to Threonine (Red). This leads to repeating of the 22 amino acids sequence 4 times in the region. B. The location of SNP in DGRP379 is at 12446453: Guanine to Cytosine, which changes Alanine to Proline (Red). The surrounding region is a poly-Q rich sequence which was determined to be conserved in *Drosophila* species by doing a BLAST search.
