## Supplementary material for "Roles of *PKCdelta* in Photoperiod and Circadian Regulations in *Drosophila melanogaster*": Figure S8

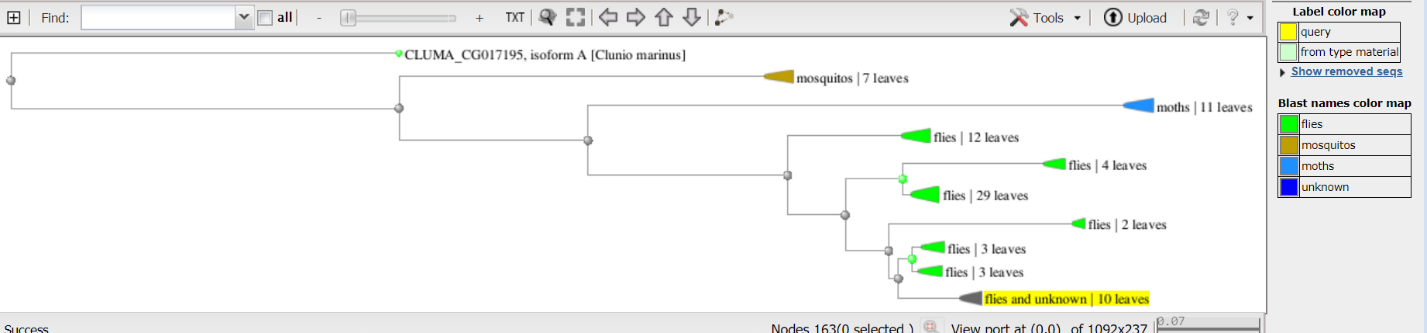

**Figure S8.** Phylogenetic analysis of *PKCdelta* RE transcript.

Tree diagram showing the evolution of the RE transcripts in *Drosophila melanogaster* (highlighted).
