## Supplementary material for "Roles of *PKCdelta* in Photoperiod and Circadian Regulations in *Drosophila melanogaster*": Figure S9

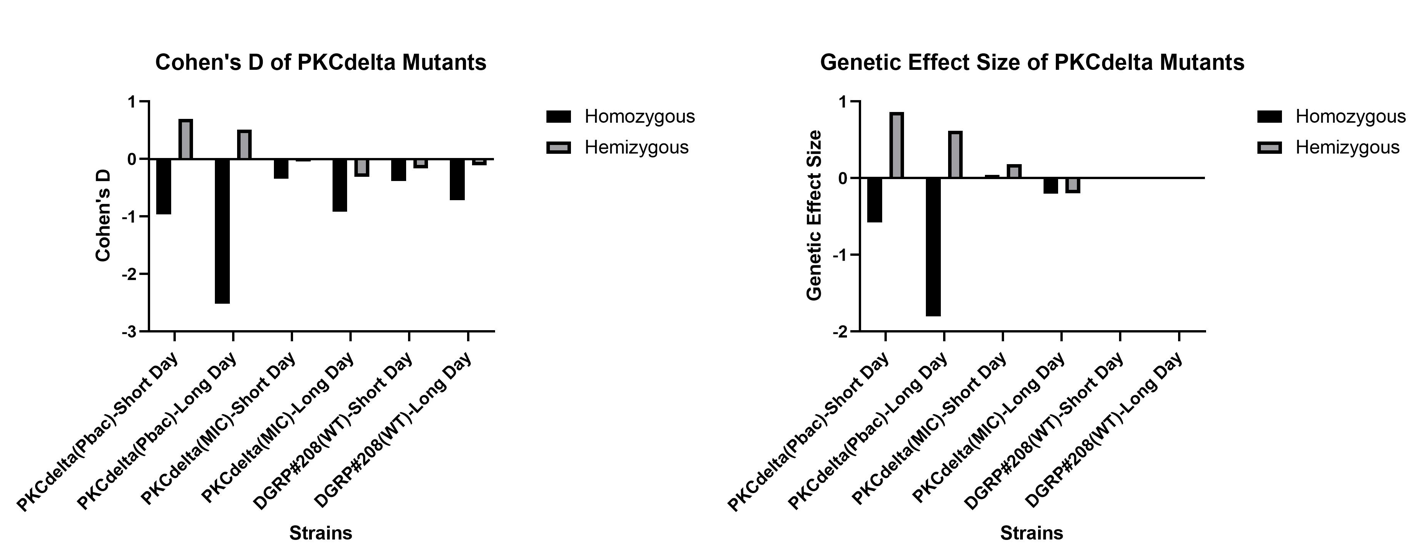

B *PKCdelta*:Pac mutants

A *PKCdelta*:Pac mutants

**Figure S9.** Cohen’s D and Genome Effect Size of *PKCdelta* mutants.

Cohen’s D of the *PKCdelta* homozygous females (Black), hemizygous (Gray) males under LD and SD compared to DGRP 208 WT. A. CD are obtained by subtracting EQ from SD and LD. There is a difference in photoperiod response between homozygous and hemizygous flies of *PKCdelta:*Pbac. This difference is reduced in the *PKCdelta:*MIC flies and WT flies. B. GES of the mutants compared to WT. *PKCdelta:*Pbac homozygous flies under SD and LD photoperiod has negative GES, while the *PKCdelta:*Pbac hemizygous flies have positive GES under SD and LD. The *PKCdelta:*Pbac homozygous and hemizygous mutants under both photoperiod condition is significantly different from the WT. *PKCdelta:*MIC mutants show a reduced photoperiod response and there is no significant difference between homozygous and hemizygous flies compared to WT.
