## Supplementary material for "Roles of *PKCdelta* in Photoperiod and Circadian Regulations in *Drosophila melanogaster*": Figure S10

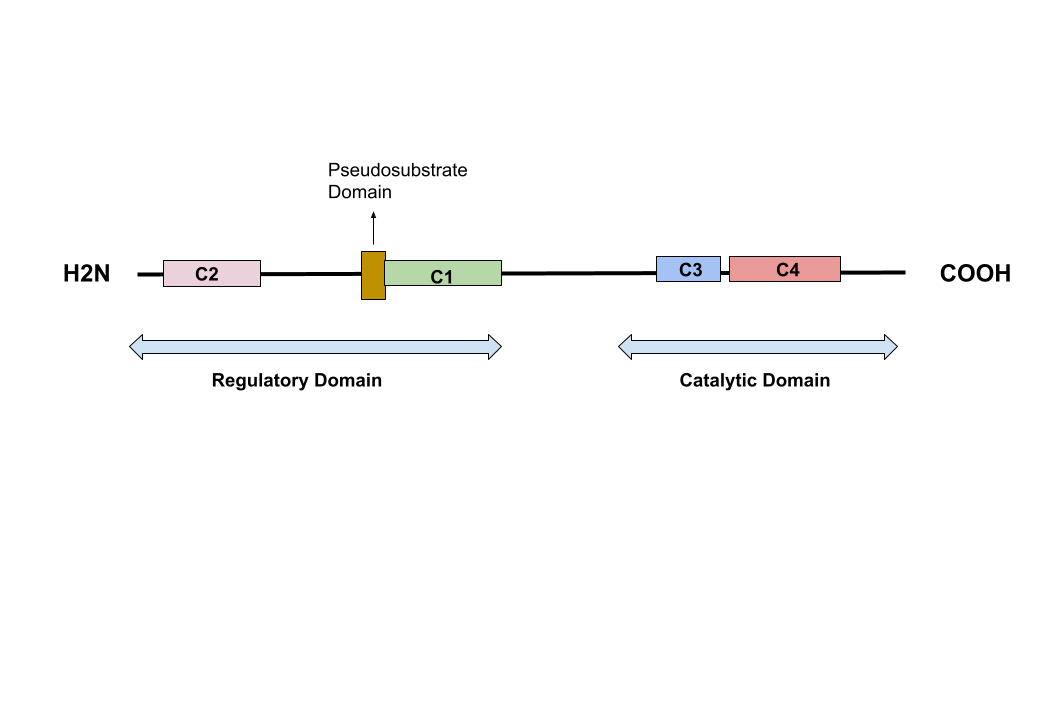

**Figure S10.** The regulatory and catalytic domains of *PKCdelta*.

Towards the C-terminal this gene contains protein kinase catalytic domain which cover exons 7-15. The catalytic domain has an ATP-binding region (C3), a catalytic active substrate binding region (C4). The N-terminal regulatory domain has an inhibitory pseudosubstrate region, and zinc-finger like sequence which is cysteine rich (C1) (Yoshida, 2007). The catalytic domain in protein kinases is known to transfer phosphate from ATP to amino acid residues which ultimately causes conformation changes in the protein (Hanks et al., 1988). Sequencing data for the mutants were deposited at NCBI, PRJNA812613.

Yoshida, K. (2007). PKCδ signaling: Mechanisms of DNA damage response and apoptosis. Cellular Signalling *19*, 892–901.

Hanks, S.K., Quinn, A.M., and Hunter, T. (1988). The protein kinase family: conserved features and deduced phylogeny of the catalytic domains. Science *241*, 42–52.
